# Sleep-Deep-Net learns sleep wake scoring from the end-user and completes each record in their style

**DOI:** 10.1101/2023.12.22.573151

**Authors:** Fumi Katsuki, Tristan J. Spratt, Ritchie E. Brown, Radhika Basheer, David S. Uygun

**Affiliations:** VA Boston Healthcare System and Harvard Medical School, Dept. of Psychiatry, West Roxbury, MA 02132, USA

**Keywords:** Automated Sleep-Wake scoring, Deep learning, Transfer learning, Translational Sleep Research, Basic Sleep Research, *In Vivo* Pharmacology, Hypnotics, Alzheimer’s Disease Model

## Abstract

Sleep-wake scoring is a time-consuming, tedious but essential component of clinical and pre-clinical sleep research. Sleep scoring is even more laborious and challenging in rodents due to the smaller EEG amplitude differences between states and the rapid state transitions which necessitate scoring in shorter epochs. Although many automated rodent sleep scoring methods exist, they do not perform as well when scoring new data sets, especially those which involve changes in the EEG/EMG profile. Thus, manual scoring by expert scorers remains the gold-standard. Here we take a different approach to this problem by using a neural network to accelerate the scoring of expert scorers. Sleep-Deep-Net (SDN) creates a bespoke deep convolution neural network model for individual electroencephalographic or local-field-potential records via transfer learning of GoogleNet, by learning from a small subset of manual scores of each EEG/LFP record as provided by the end-user. SDN then automates scoring of the remainder of the EEG/LFP record. A novel REM scoring correction procedure further enhanced accuracy. SDN reliably scores EEG and LFP data and retains sleep-wake architecture in wild-type mice, in sleep induced by the hypnotic zolpidem, in a mouse model of Alzheimer’s disease and in a genetic knock-down study, when compared to manual scoring. SDN reduced manual scoring time to 1/12. Since SDN uses transfer learning on each independent recording, it is not biased by previously scored existing data sets. Thus, we find SDN performs well when used on signals altered by a drug, disease model or genetic modification.

**STATEMENT OF SIGNIFICANCE:** Sleep medicine is often critically advanced by translational research based on *in vivo* electrophysiologic mouse data. A necessary but time-consuming step in this field is scoring epochs of recordings into wakefulness, non-rapid-eye-movement sleep and non-rapid-eye-movement sleep. Despite efforts to automate this, manual scoring remains the gold-standard since automatic methods poorly handle data that is not similar enough to data used during development. Here, we describe a novel automated sleep scoring method that involves retraining a deep-convolution-neural-net capable of computer vision to score sleep-wake patterns after learning from a small set of manual scores within a record. This avoids biasing the model to expect data to be the same as its training set from previous records.

## INTRODUCTION

Assigning behavioral states into wakefulness, non-REM sleep or REM sleep has been an essential component of clinical and pre-clinical sleep research ever since the first all-night recordings of sleep in humans.^1^ So called sleep-wake scoring is a necessary step in the workflow from electroencephalographic (EEG) or local-field-potential (LFP) data collection to experimental observations in basic and translational sleep research, allowing analysis of the amount of sleep states, sleep architecture and alterations in specific sleep oscillations. However, manual sleep-wake scoring requires concentration, is time consuming, labor intensive and laborious. New expert scorers require training from experienced scorers. Scoring data from rodents has unique challenges to scoring data from humans due to the smaller separation of EEG amplitude differences between states^2^ and the need for scoring shorter epochs which are conventionally 4 or 10 seconds^3, 4^, contrasting with 30 second epochs^5^, as standard in scoring data from humans. Moreover, in animals there exists no professional standard handbook for reference, as used to score data from humans.^6, 7^ Instead, criteria for scoring rodent data come from methods sections of research articles, typically described in a single paragraph.^3^ Since subsequent automated analytic methods can be performed rapidly on the EEG/LFP records once the sleep-wake record is scored, the manual scoring step can be considered a bottleneck. These considerable limitations of manual scoring have created a strong impetus to automate sleep-wake scoring, with efforts dating back decades.^8^ Older approaches may work relatively well for clearly defined criteria of wake, rapid-eye-movement sleep (REM) & non-REM (NREM), typical of wild-type (WT) mouse data. However, these methods typically work less well in studies which require scoring mouse EEG/LFP/electromyographic(EMG) data involving an experimental intervention, such as a genetic modification, disease model treatment, or drug. These experimental interventions may alter the quality of sleep, as defined by the EEG waveforms, altering the definition of the sleep-wake state. Thus, manual scoring by human scorers has remained the gold-standard in the field.^8^

Recently, machine learning based algorithms have been developed to address the problem of automating sleep wake scoring. In the last 4 years, at least 7 machine learning based sleep-wake scoring algorithms have been published. These algorithms work by creating a novel machine learning model that is trained on a data set. A major strategy used to increase performance, is to provide the largest data set available for training the network. These models are then required to score new data based on what it has learned from the data with which they are familiar.^9–16^ However, one concern with this approach is that networks can often perform poorly when they are asked to classify new data that they have not encountered before.^17^ This could be the case with data from a different laboratory or unfamiliar experimental treatments, such as alterations in EEG/EMG produced by pharmacological agents, genetic manipulations, and/or disease models.

We decided to take a novel approach to this problem. Instead of building a new unique machine learning model trained on data and scores from our laboratory, we employ transfer learning to considerably speed up the scoring of data by an expert scorer. This is a deep-learning based technique that leverages a pre-trained highly sophisticated deep neural network already capable of computer vision, and retrains it at a superficial level to score images that represent NREM, REM and wakefulness. For this we utilized GoogleNet, winner of the 2014 ImageNet challenge, a pre-trained deep neural network.^18^ Our new method, which we term Sleep-Deep-Net (SDN), creates a newly tailored model for each individual file via transfer learning of GoogleNet, which is retrained on a small set of manual scores from that file provided by the end user. Since learning is based on a small subset of manual scores performed on each recording of a mouse’s EEG/LFP/EMG signal, SDN automation remains faithful to the learned EEG/LFP/EMG pattern of each mouse. SDN is not familiar with older existing mouse EEG/LFP/EMG data from our, or any other, labs. Thus, when a new investigator uses SDN, it learns to score from them. Similar approaches have been used in human data and wearable technology.^19, 20^ However, this approach has not been applied to data from mice, the most commonly used animal model in pre-clinical sleep research, as far as we are aware. It is important to note that no automated sleep scoring method works perfectly on every file. Especially, scoring REM appears to be one of the major challenges for the automated scoring methods. An automatic indicator of poorly scored REM would be valuable in determining whether to use a particular set of automated scores. With SDN, we developed a way to identify files yielding the lowest REM reliability, which provides the users options to either exclude or manually score those files. No other method does this. We aim to make this freely available to the sleep research community to accelerate basic and translational research in mice.

## METHODS

### Mouse sleep-wake datasets and surgical procedures

Two mouse models with EEG and one disease mouse model with local field potential (LFP) electrode recordings were used for developing and validating the SDN automated scoring. For EEG/EMG recordings, we used data sets previously obtained from mice expressing the enzyme, Cas9, in parvalbumin neurons (PV-Cas9 mice), before and after the CRISPR knockdown of the GABA_A_ receptor α3 subunit in the Thalamic reticular nucleus.^21, 22^ For pharmacology work, male C57BL/6 mice (Strain #000664 Jackson Laboratory) were used. As in our previous published work^21, 22^, these mice were implanted with frontal cortex (AP +1.9 mm, ML ±1.5 relative to bregma) EEG screw electrodes (Pinnacle Technology Inc.; Kansas, United States; Part # 8403) with a reference screw electrode above the cerebellum (AP bregma −6 mm ML = 0) and a ground electrode screw at bregma AP −3 mm, ML +2.7. EMG electrodes were positioned under neck muscles. For validations using LFP/EMG data we used transgenic mice which were produced by crossing PV-Cre mice (Stock #: 017320; Jackson Laboratory) and APP/PS1 mice, a well-known β-Amyloid overexpressing model of Alzheimer’s disease (Stock #: 034829-JAX; MMRRC stock #34829). These PV-cre/APP/PS1 mice were used for the LFP recordings in which the tungsten electrodes were implanted in the prefrontal cortex (AP +1.9 mm, ML –0.45 mm, DV –1.7 mm) with a reference and a ground screw electrode above the cerebellum (AP bregma −5.8 mm, ML ±1.25). In all surgeries, EEG or LFP, attached wires were soldered to Pinnacle Technolology Inc. headmounts (Part # 8201-SS) which were then secured in place by Keystone industries Bosworth Fastray (Part # 0921378) dental cement. Mice were allowed at least one week to recover before recordings were made. Surgeries were performed on a Leica stereotaxic system under isoflurane (1.5– 4% in O2) anesthesia monitored by breathing rate, pedal withdrawal and tail pinch reflexes. Meloxicam (5 mg/kg; intraperitoneal or subcutaneous) was given immediately at the end of surgery and again 22–24 h later, to treat post-surgery pain.

### Sleep-wake recording procedures

We recorded EEG/LFP and EMG on Pinnacle Technology Inc. three-channel (2 EEG and 1 EMG) mouse systems (Part # 8200-K1-SL), via native software (Sirenia Acquisition). Mice were habituated by tethering them to their system via pre-amplifiers for mice (Pinnacle Technology Inc. Part # 8202-SL) for 24+ hours. 24-hour recordings were collected between zeitgeber time (ZT) 0 – 24. Data were sampled at 2 kHz, amplified 100x and low pass filtered at 600 Hz. Zolpidem tartrate was administered 5mg/kg, i.p., at ZT 6. All experiments were approved by the Institutional Animal Care and Use Committee of VA Boston Healthcare System and conformed to National Institute of Health, Veterans Administration and Harvard Medical School guidelines.

### Manual sleep-wake scoring

We used Sirenia Sleep (version 2.0.1) to manually score a 2-hour section of 24hour EEG/EMG records focusing on parts of the sleep-wake recording (ZT 2.5 – 4.5) where we could identify as including REM, since REM is much less abundant than wake and NREM. For the pharmacology work, we manually scored a 20 minute period spanning and roughly centered on the injection of zolpidem. Zolpidem is short acting^23^ so we focused on a 3 hour period for analysis and hence we also manually scored a shorter (20 min) period for SDN training. Manual scoring was performed as previously described^21^, using 4 s epochs. EEG/EMG and LFP/EMG signals were exported as EDFs using Sirenia’s native export options. All epochs, including those scored, manually were exported as tsv files and converted to txt files. Briefly, our method for manual scoring was as follows. Epochs with a desynchronized low amplitude EEG signal and a large EMG signal that did not necessarily need to appear phasic were labelled wake. Epochs containing mostly large amplitude, slower waves, and a low amplitude EMG signal – with the exception of brief bursts suggesting a twitch – were labeled NREM. Epochs with a repetitive stereotyped ‘sawtooth’ EEG signal oscillating at 5–9 Hz (theta), with a very flat EMG signal were labeled REM.

### Wavelet transform of signals

First, for each manually scored epoch, continuous wavelet transforms were performed using the Matlab cwtfilterbank function with 32 voices per octave on three-epoch wide sections of raw EEG/EMG or LFP/EMG – the score of the middle epoch was assigned to each section, flanked by its neighboring epochs. The EEG/EMG or LFP/EMG wavelet transforms were converted to RGB images, with the top half showing the EEG or LFP wavelet transforms and the bottom half showing EMG wavelet transforms per epoch-triplate, in the jet color scheme. Images were sized to 224×224 and saved as jpegs. The jpeg files were then saved in folders, each labeled with the following categories: “W”, for wake, “N”, for NREM; “R”, for REM. The same conversion to EEG/EMG or LFP/EMG wavelet transform image files was performed for every epoch in the EEG/EMG or LFP/EMG record and saved in a folder to be classified by the algorithm, once trained. We wrote and modified code to perform the wavelet transform of signals by consulting Matlab documentation.^24^

### Retraining GoogLeNet for each EEG/EMG or LFP/EMG record

We performed transfer learning of GoogLeNet using the Matlab Deep learning Toolbox. GoogLeNet was obtained from Matlab’s pretrained deep neural nets. As it exists without modification, GoogLeNet classifies images into one of 1000 object categories, such as a mouse, keyboard or pencil etc.^25^ We retrained GoogLeNet to classify images representing wake, NREM & REM – jpegs of wavelet transforms described above. In a neural net, so called convolutional layers extract features, which the final learnable layer and classification layers use to classify the images.^24^ In GoogLeNet, “loss3-classifier” and “output” are these final learnable and classification layers, respectively, containing the information about how features are combined to create class probabilities and predictions^24^ – I.E. sleep-wake scores once it is retrained. Prior to our modification: “loss3-classifier” corresponds to the 1000 categories pertaining to object recognition (mouse, keyboard or pencil etc). This “loss3-classifier” layer was replaced with a new layer that instead has the same number of categories as sleep-wake states present in the manual scores (wake, NREM, REM). Of note, users must manually score some of each state that is needed. For example, a user should not score a period of data devoid of, or poorly representing, REM. The output layer was also replaced to reflect the number of sleep-wake states represented in the manual scores. We replaced a ‘dropoutLayer’, with one that has a dropout probability of 0.6 to prevent over-fitting.^26^ Otherwise, the GoogLeNet architecture was unchanged. Using these modifications to GoogLeNet, novel deep-nets were trained, per file, using the Matlab ‘trainNetwork’ function. We wrote and modified code to retrain GoogLeNet by consulting Matlab documentation.^24^ The images assigned for validation were 20% of the images generated from each class (sleep-wake state), selected at random using the splitEachLabel function. we selected dropout probability based on Matlab documentation^24^, Maximum Epochs were determined empirically with our publicly available data set.^22^ We chose 10 epochs after initial trials with the default setting of 20 epochs. We observed validation accuracy and loss plateaued by 10 epochs in all files, and F1 scores improved. 5 epochs led to lower F1 scores so we used 10 epochs for the rest of the study. We also tested other mini-batch sizes, weight learn-rates and bias learn-rate factors, but these did not improve performance. Therefore, these and other user defined settings were set to default as follows: ‘Stochastic gradient descent with momentum’, ‘Mini-Batch Size’ = 15, ‘Initial Learn Rate’ = 1e-4, ‘Validation Data’ = images assigned for validation, ‘Validation Frequency’ = 10, ‘Verbose’ = 1. To increase speed, we used the GPU as the ‘Execution Environment’. Finally, a newly re-trained model tailored to each file was used to classify the non-scored epochs for each file using the Matlab classify function.

### REM Quality Control to identify files that could not be scored reliably by SDN

In order to further improve accuracy of REM scoring, we developed the post-hoc corrections which involves a few extra steps. We discovered that the Matlab “classify” function does not simply assign the class based on its highest certainty estimate (one of the classify functions output variables). In other words, some REM epochs are assigned with a higher certainty estimate for wake or NREM. First, we re-labeled each REM epoch based on its highest certainty value. Next, we corrected cases where REM immediately following wake which is an abnormal state transition that could only been observed in the specific cases such as the mouse models of narcolepsy.^27, 28^ We first found the Wake to REM cases and identified the surrounding scores for each case. If a wake→REM case was preceded by more than 3 epochs of wake, then the REM was replaced with wake, assuming that an epoch within a wake bout was incorrectly scored as REM. If a Wake to REM case was followed by more REM epochs and had less than 5 wake epochs prior to it, then epochs between the last NREM and the wake to REM case were replaced with REM, assuming that the epochs at the transition from NREM to REM were incorrectly scored as wake. Our post-hoc correction of REM scores effectively removed REM from files that could not be scored accurately. This feature lets us identify the files that could not be reliably scored by SDN. For the PV-Cas9 mice dataset, only three out of fourteen files used were flagged as unreliable. In this example, these three files could then be scored manually and used in a study. For simplicity, we removed those three from the validation data shown in the results section.

### Secondary optional post-hoc correction of REM epochs

In addition to the post-hoc corrections of REM described above, we wrote a custom function termed ‘rule_123’ which first searches for REM epochs preceded by a single NREM epoch, then replaces the REM and the following epoch as NREM. Next, rule_123 searches for short REM bouts – from a single epoch up to four epochs – which we did using 4s epochs – and relabels them as wakefulness.

### Hardware and software specifications

Work was performed on either a computer with an Intel(R) Core(TM) i7-8700 (clock speed: 3.20GHz), 32.0 GB RAM, and a NVIDIA GeForce RTX 2080 GPU or an Intel(R) Core(TM) i9-11900F (clock speed: 2.50GHz, 128 GB RAM, and a NVIDEA GeForce RTX 3090 GPU. Both ran Windows 10 Pro and MATLAB R2023b. Using a GPU with specifications similar to the ones listed here is highly recommended since it speeds up the computationally demanding processes such as the wavelet transform of signals and the network retraining by ∼7 times faster than using CPU only.

### The time-frequency spectrogram

The time-frequency spectrogram was produced using the multi-taper method (Chronux Toolbox; Chronux.org) function^29^ (5 tapers with 10 s sliding window in 100 ms steps).

### Statistics

Comparisons were made by either, Pearson’s correlations, two-way ANOVAs or t-tests as described. F1 scores were calculated as the harmonic mean of precision and recall, where precision was the fraction of detections of a class that were correct; recall was the fraction of correct classes that were detected.

## RESULTS

To assess how reliable our transfer learning-based algorithm, SDN, was at classifying scores after training on manually labeled scores, we first calculated F1 scores for the three vigilance states typically tabulated in mouse EEG and overall. Automated scores by SDN provided high F1 scores (**Figure 1A**) for wakefulness (0.97±0.003), NREM (0.95±0.004), REM (0.86±0.011) and overall (0.96±0.004) in eleven 24-hour EEG/EMG records. Importantly, the proportion of each vigilance stage is similar when EEG/EMG records were scored by SDN vs a human (**Figure 1B&C**). A two-way ANOVA revealed significance only in the ‘sleep-wake state’ factor [F (2,60) = 2659, p < 0.001], with no significance in the ‘score-method’ factor [F (1,60) = 0.01, p = 0.91] or interaction [F (2,60) = 1.82, p = 0.17]. Moreover, the proportions of states determined from automated scores vs manual scores correlated very highly in wakefulness (**Figure 1D**, r = 0.94, *p* = 0.000007), NREM (**Figure 1E**, r = 0.85, *p* = 0.0005), and REM (**Figure 1F**, r = 0.87, *p* = 0.0002), as determined by Pearson’s correlation. For this we used 11 separate 24-hour records from six different mice. To address whether these correlations were significant by virtue of the within animal replicates, we performed additional correlations using only a single record per mouse and still found significant correlations for wake (ρ = 0.95, *p* = 0.002), NREM (ρ = 0.92, *p* = 0.005) and REM (ρ = 0.85, *p* = 0.02). Of note, the 11 files shown here were a subset of 14 files from seven mice. After utilizing our “REM Quality Control” (see methods), we were able to identify 3 out of 14 files as unreliably scored, which we determined as described next.

**Figure 1.**
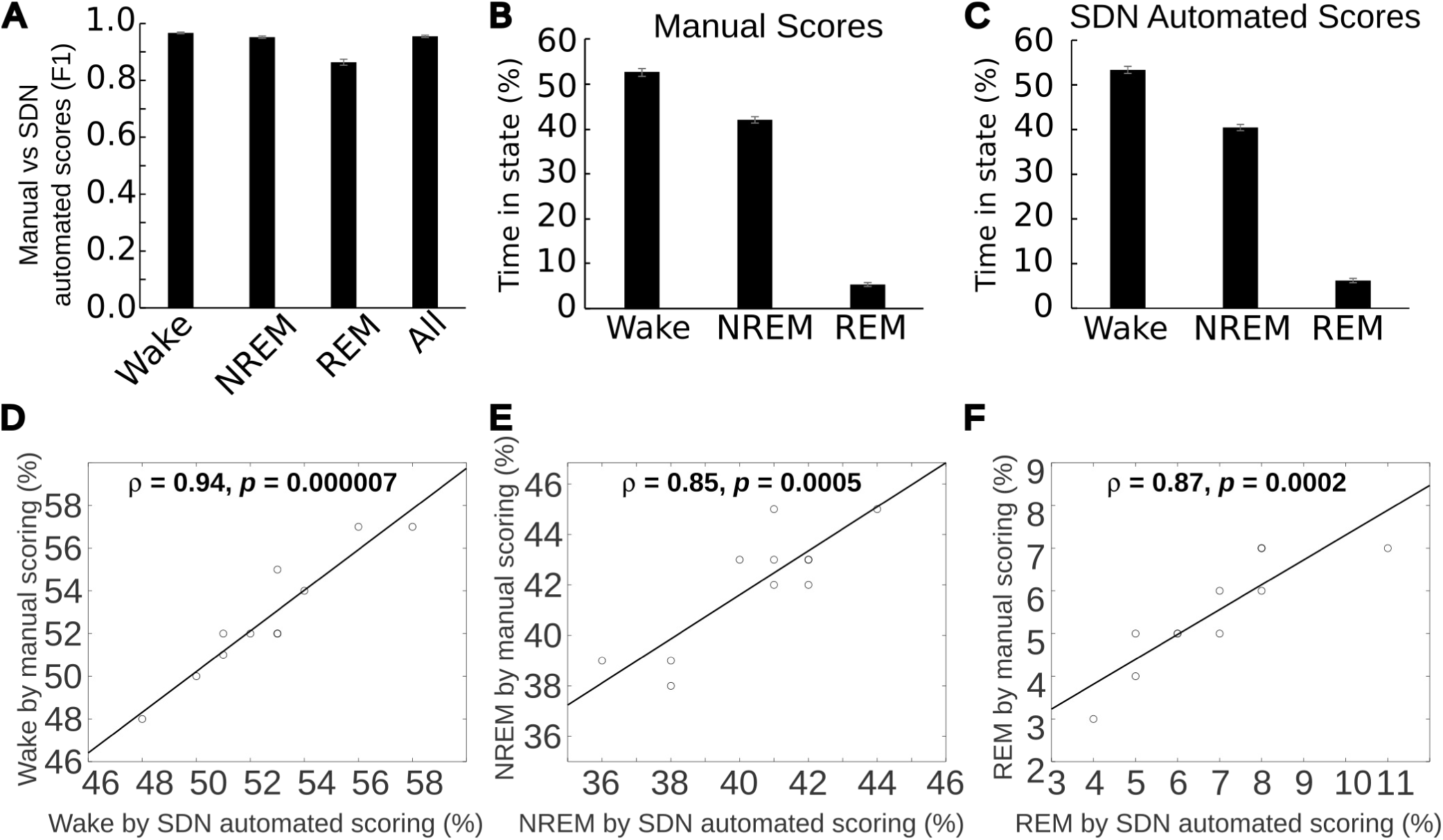
Sleep-Deep-Net (SDN) scores mouse EEG/EMG signal with high reliability compared with human expert scores and retains the proportion of sleep-wake states. **A.** F1 scores for wakefulness, NREM and REM reveal reliable scoring respective to manual scores by the expert scorer on which SDN was trained. **B.** Shows the proportion of wake, NREM and REM from 24-hour records when determined by manual scoring. **C.** Shows the proportion of wake, NREM and REM from 24-hour records when determined by SDN automated scoring. On a 24-hour record-by-record basis, the proportion of wakefulness (**D**), NREM sleep (**E**), and REM (**F**) correlate highly when determined by manual scoring vs SDN automated scoring. Data were from eleven 24-hour recordings from six mice.

Initially, we noticed that REM F1 scores were lower than those of Wake and NREM. We were interested in developing an automated “REM Quality Control” procedure that would reliably indicate EEG/EMG records in which SDN REM scores were unreliable. We thought this would be useful because investigators using SDN would not be able to calculate F1 scores on new unscored data. Unreliable EEG/LFP/EMG records could then be scored manually, or excluded from further analysis. First, we looked at a variable called “validation accuracy”, provided by Matlab during training. But this was a poor correlate of final REM F1 scores (**Figure S1A**, r = 0.03, *p* = 0.46). However, using a simple “REM Quality Control” procedure, we were able to slightly improve F1 scores in most of the files. Importantly however, this had the effect of drastically lowering REM F1 scores that were already the lowest (**Figure SB, red**). These corrected REM scores yielded F1 values that became correlated with the validation accuracy value (ρ = 0.54, *p* = 0.02). Unfortunately, two files with validation accuracy in the same lower range still produced high REM F1 scores (**Figure S1B, orange filled**), which would lead an investigator to unnecessarily score two files manually. However, we found our REM Quality Control procedure decimated REM selectively in unreliably scored files, flagging only the three out of fourteen files from our full data set (**Figure S1C, red**). The final proportion of REM sleep in each file became an excellent indicator of F1 scores (**Figure S1C**), correlating very highly (ρ = 0.81, *p* = 0.0002). In a typical experimental scenario, these three files could then be scored manually and used in a study. For simplicity, we removed those three from this validation. Unless mouse models used in a study are known for the direct wake to REM transition, we recommend all SDN users employ this post-hoc correction, as it was necessary to identify files yielding poor reliability of REM scoring.

Finally, we made a secondary optional post-hoc correction to REM scores which further improved F1-scores. This optional post-hoc correction basically removed REMs that was preceded by only one epoch of NREM or had a very short bout length. Before developing rule_123, we had observed that recall for REM (0.92±0.01) was considerably higher than precision (0.82±0.02). Thus, if we could identify major sources of false positives, we could relabel them as another state. We observed that the algorithm commonly scores REM following short bouts of NREM, though manual scores produce these instances very rarely (**Figure S1A**). Therefore we employed this simple strategy to identify a triple-epoch sequence of “wake or REM”→NREM→REM to identify REM preceded by a single NREM epoch in the SDN scores. We observed that in most of these epochs, manual scoring had identified the “REM” epoch as NREM. So, we relabeled that SDN scored REM as NREM. Next, we observed that SDN led to more time-weighted REM bouts in the shortest bin at the expense of the next shortest bin (**Figure S1B**). We found relabeling very short REM bouts as wake improved REM F1 without reducing the wakefulness F1 score from 0.97 and corrected the profile of time-weighted REM bouts (**Figure 2B&C, right**). We determined the minimum REM epoch duration by increasing it one epoch at a time up to four epochs. We found REM F1 scores increased by 0.1 per epoch and plateaued at three REM epochs. We propose this custom Matlab function we term ‘rule_123’ as an optional feature investigators may choose to use based on comparisons of F1 scores and time-weighted bout analysis in their own data sets previously scored manually, then use those optional settings with novel data sets. We found it worked well with our EEG data.

**Figure 2.**
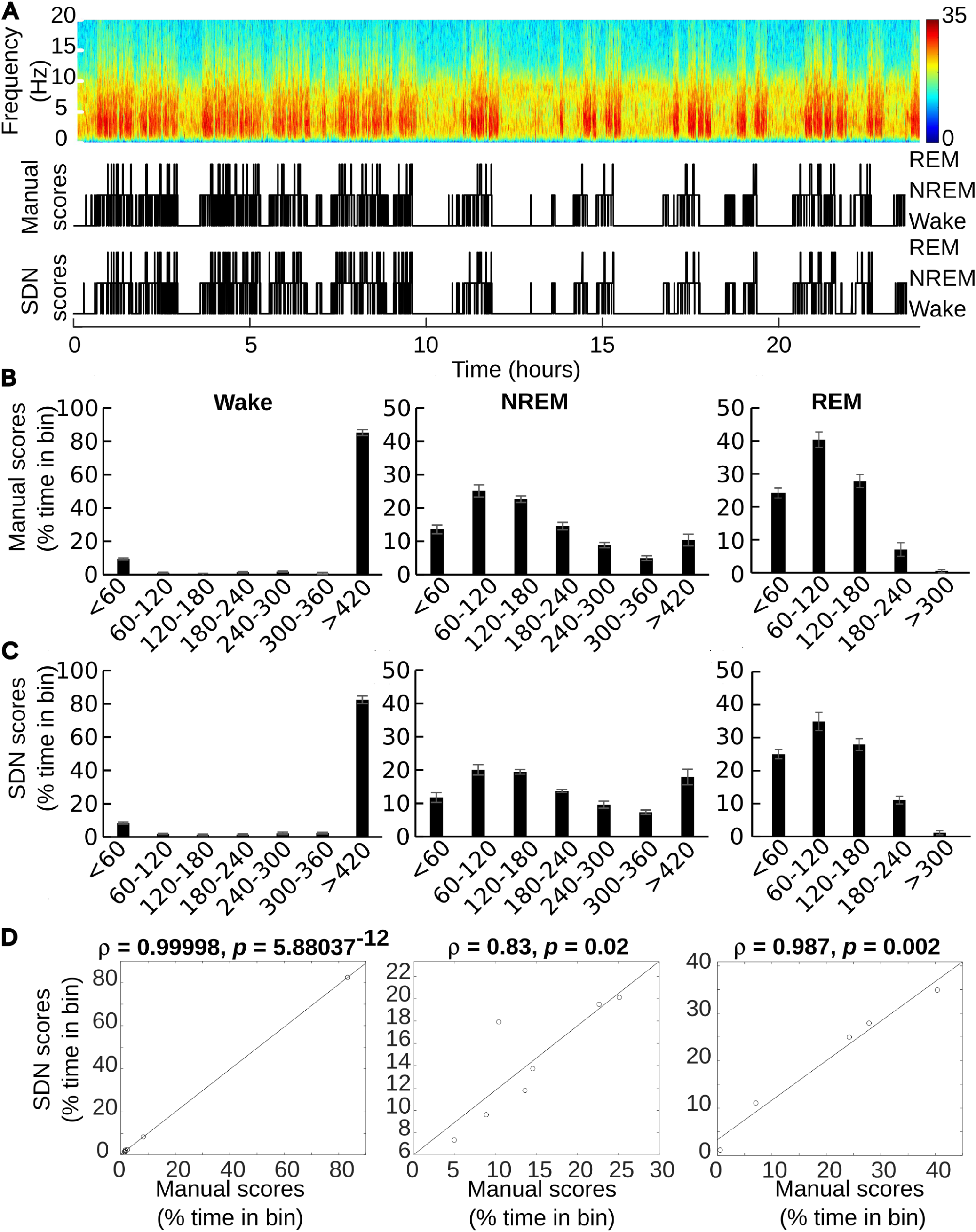
Sleep-Deep-Net (SDN) retains sleep architecture of manual scoring. **A.** A representative multitaper time-frequency plot of EEG from a 24 hour record with aligned hypnograms drawn from manual scores and SDN automated scores. **B.** Time-weighted bouts of wakefulness, NREM and REM as determined from manual scores. **C.** Time-weighted bouts of wakefulness, NREM and REM as determined from ADN automated scores. **D**. Scatter plots with fitted least squares line reveal close relationship between time-weighted bouts for all three sleep-wake states on a record-by-record basis. Pearson’s correlation coefficients reveal statistically highly significant correlations between all three sleep-wake states comparing manual vs SDN automated scores. Data were from eleven 24-hour recordings from six mice.

Though high F1 scores are needed to indicate widespread agreement overall and within each state, a deep analysis of sleep-wake architecture was necessary to determine whether SDN would produce scores leading to results similar to those of an expert scorer, if we were to use the method in real research applications. Next, we evaluated sleep architecture in the EEG/EMG records when scored by SDN vs manually. **Figure 2A** shows a representative time-frequency plot of EEG with aligned hypnograms comparing manual scores with SDN scores to illustrate close alignment across the entire 24-hour EEG/EMG record. Thus, SDN generated scores retain the sleep-wake architecture of manual scoring. To perform fine-grained evaluation of sleep-wake dynamics, we performed bout analysis. For this, we analyzed time-weighted bouts as previously described first by Mochizuki *et al.*^30^ and used later by our group.^21^ We found the profiles of time weighted bouts for wake, NREM and REM as scored manually (**Figure 2B**) and the profiles of time weighted bouts for each state as scored by SDN (**Figure 2C**) correlated very highly for all three states (**Figure 2D**): Wake (ρ = 0.99998, *p* = 5.88037^-12^), NREM (ρ = 0.83, *p* = 0.02) and REM (ρ = 0.987, *p* = 0.002).

Next, we wanted to test the performance of SDN under a condition known to affect the quality of sleep by altering NREM EEG waveforms in mice^23, 31^ to test the versatility of SDN with data other than WT EEG/EMG signals. We recorded three hours of EEG/EMG from four mice and delivered a hypnotic dose of zolpidem at 1 hour after the start of the recording. **Figure 4A** shows a representative time-frequency plot of the EEG in one mouse preceding and following the injection of zolpidem, with aligned hypnograms from manual scores and scores by SDN. Zolpidem causes rapid onset NREM sleep that presents with increased low frequency power compared to wake but reduced broad band frequency power compared to NREM. Qualitatively, the sleep architecture as defined by manual scores is conserved by the automated scores generated by SDN. SDN produced F1 values in the same range as in WT data for NREM and REM, producing the highest F1 for NREM. SDN performed worse than WT data for wakefulness. Despite poorer performance in wake, this result supports utility of SDN in studies of hypnotics studies since the main therapeutic effect of zolpidem is to induce NREM sleep. Thus, EEG signals of NREM induced by zolpidem could be used to study the profile of the drug. Moreover, though epoch to epoch agreement of wake is lower than preferred, the proportion of states within the recording is preserved fairly well (**Figure 4C&D**) with a percent change to wake of only 26.08%(±13.23), NREM only −6.86%(±2.74) and REM 0.00%(±13.61).

**Figure 3.**
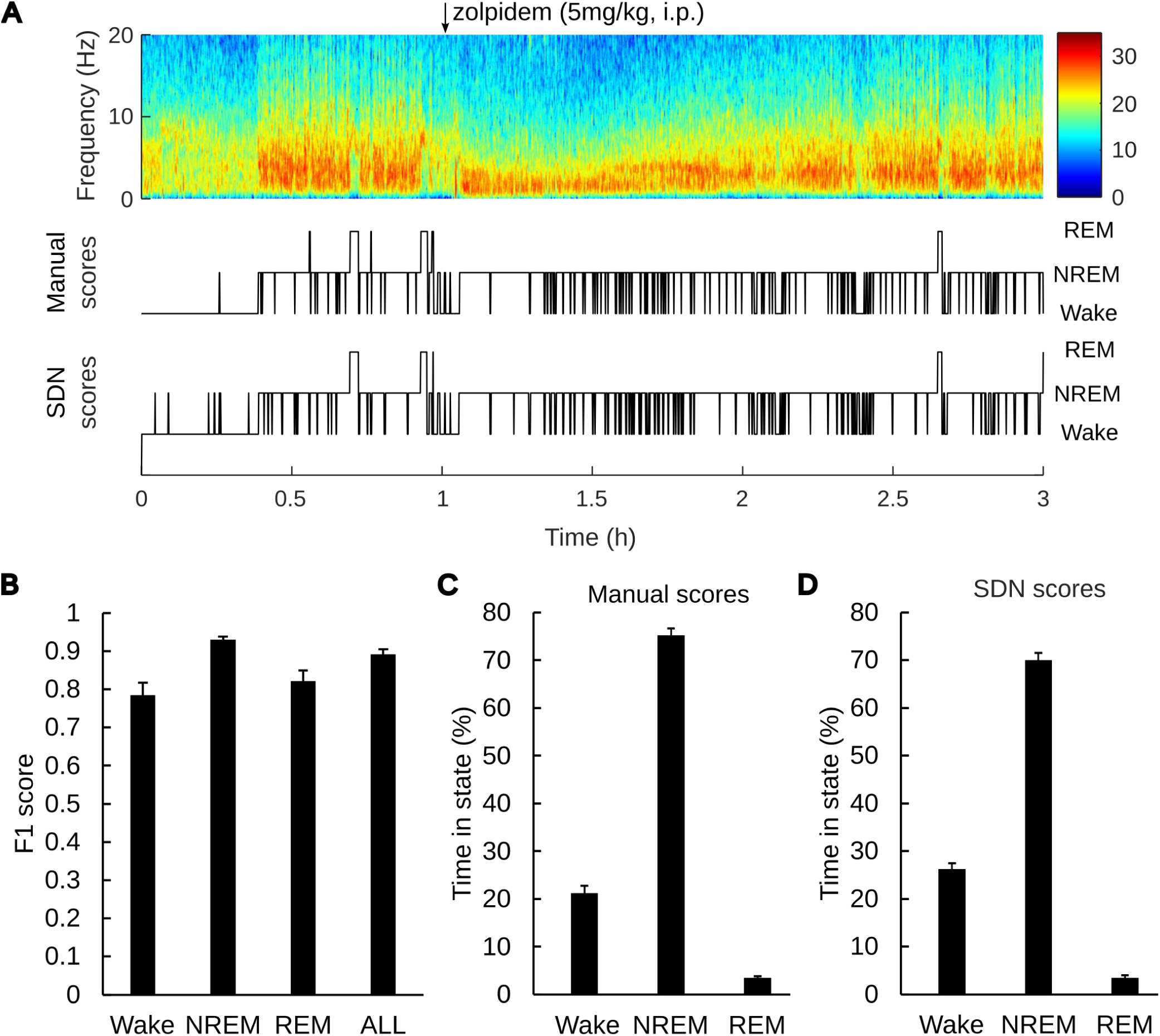
Sleep-Deep-Net (SDN) retains sleep architecture of hypnotic-drug (zolpidem) induced sleep compared with manual scoring. **A.** A representative multitaper time-frequency plot of EEG from a 3-hour record in which 5 mg/kg zolpidem was delivered by intraperitoneal injection with aligned hypnograms drawn from manual scores and SDN automated scores. **B.** F1 scores reveal high reliability of drug induced NREM sleep and overall reliability, with reliability of REM in a similar range to WT records. The relative time spent in each sleep-wake state as determined by manual scores (**C**) vs SDN scores (**D**) shows similar values. N=4

**Figure 4.**
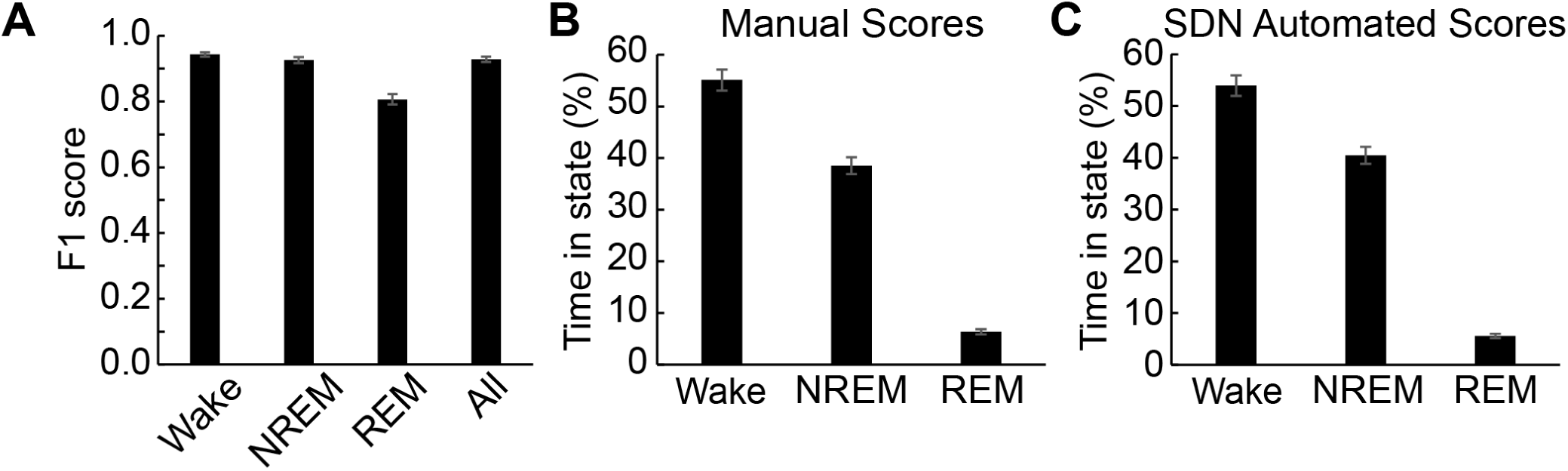
Sleep-Deep-Net (SDN) retains sleep architecture in local field potential (LFP) data from a mouse model of Alzheimer’s disease (APP/PS1) compared with manual scoring. **A.** F1 scores between manual and SDN automated sleep scores were computed with LFP recordings performed on 5 APP/PS1 mice. Each sleep-wake state and all states reveals high reliability of NREM sleep, wake, and all states combined, with reliability of REM in a similar range to the WT case shown in Fig.1. The relative time spent in each sleep-wake state as determined by manual scores (**B**) was highly similar to SDN scores (**C**). N = 5.

To further test the versatility of SDN with data other than WT EEG/EMG signals we recorded LFP from frontal cortex in an Alzheimer’s disease mouse model^32–34^ crossed with a Cre line. We used five mice that were six months old, at this age symptoms including altered sleep-wake and EEG are apparent.^35^ **Figure 5A** shows high F1 scores overall, including NREM and Wake, with slightly lower F1 scores for REM in a similar range to WT EEG data. Additionally, the relative proportion of each state is well conserved when scoring by SDN, compared with manual scores (**Figure 5B&C**). We found the proportion of each vigilance stage is similar when LFP/EMG records were scored by SDN vs a human (**Figure 5B&C**). A two-way ANOVA revealed significance only in the ‘sleep-wake state’ factor [F (2,26) = 744.83, p < 0.001], with no significance in the ‘score-method’ factor [F (1,26) = 3.91^-7^, p = 0.9995]. We found rule_123 did not improve F1 values in the LFP dataset, so it was not used here.

**Figure 5.**
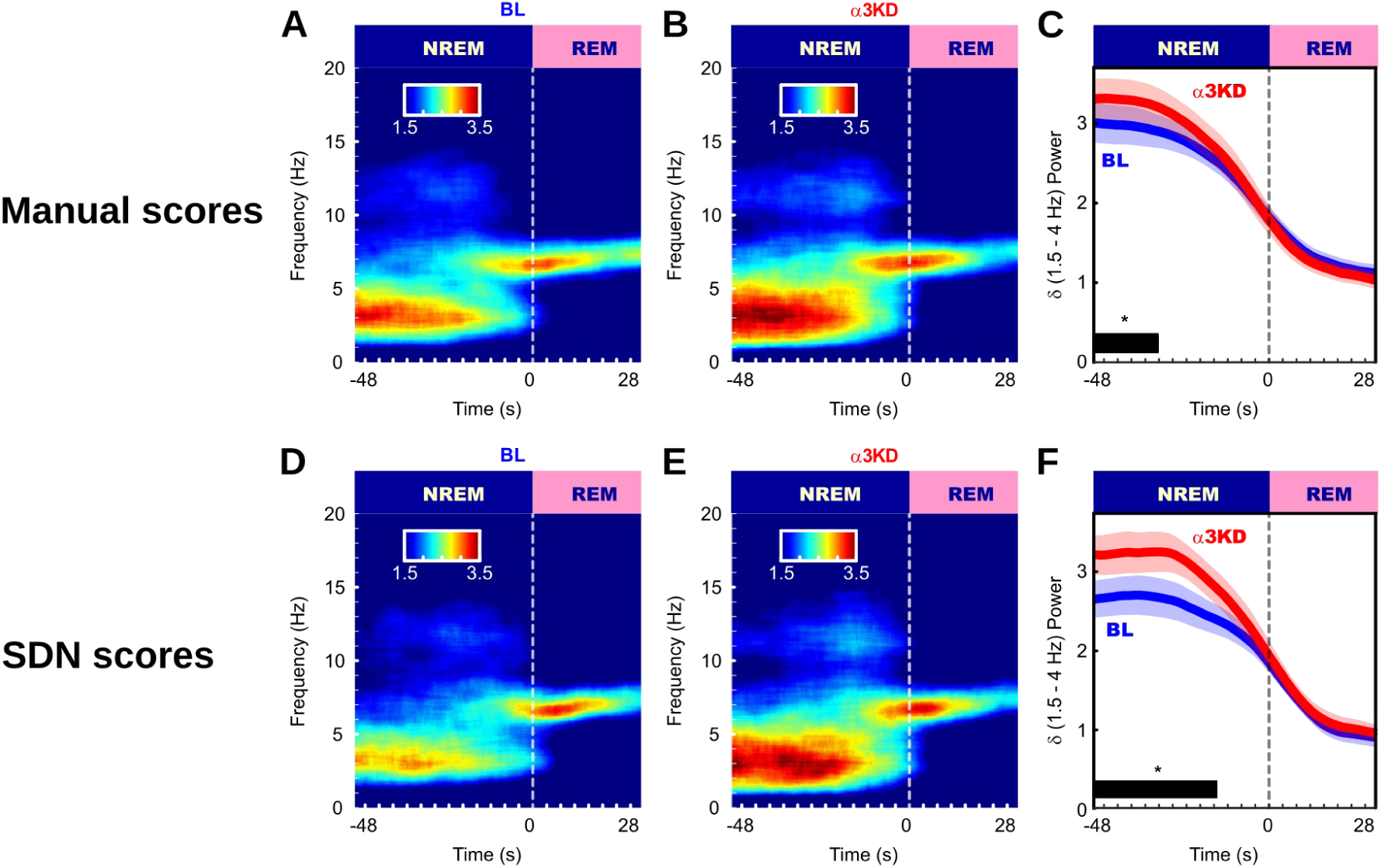
The significant increase in NREM delta power due to knocking down the expression of GABA_A_ alpha3 subunits in the thalamic reticular nucleus (Uygun et al., 2022) is reproduced when SDN scored the data. Sleep-wake states were determined by manual scoring in panels A-C, from Uygun et al., 2022. **A.** Baseline time-frequency power dynamics reveal elevated delta (1.5 – 4 Hz) during NREM prior to rapid eye movement sleep (REM) transitions, during the whole 12 h light period. **B**. Following the knock-down of alpha3 containing GABA_A_ receptors in the thalamic reticular nucleus (α3KD), high delta in NREM prior to REM transitions was further increased. **C.** In contrast to their own BL levels (blue), α3KD mice (red) produced more delta power during NREM prior to REM transitions [t *(5)* = 2.14, *p* = 0.04]. Sleep-wake states determined by SDN scoring in panels **D-F** reproduced the effects found by manual scoring [*t* (5) = 6.6, *p* = 0.0006]. Significance was tested using one-tailed paired *t*-tests. Thick lines indicate mean; envelopes indicate SEM. N=6.

Finally, we wanted to reproduce findings involving changes in NREM sleep oscillations from data previously scored manually. We recently used mouse EEG combined with *in vivo* CRISPR-Cas9 knock-down of α3 containing GABA_A_ receptors in the thalamic reticular nucleus to reveal deepened sleep as measured by elevated delta wave power enriched at NREM-REM transitions.^21^ We wanted to test whether we would have found this result had we been using SDN. We utilized SDN to re-score the data from mice that had their synaptic GABA_A_ receptors knocked-down selectively in the thalamic reticular nucleus, termed α3KD, and performed the analysis of our previous publication. **Figure 6** shows graphs redrawn from our previous publication using manual scoring showing “heightened delta power associated with α3KD was only evident during NREM sleep preceding transitions to REM sleep (**Figure 6 A-C**) [means (standard error (SEM): BL = 2.98 (0.24), α3KD = 3.3 (0.26); 11.5% change for all transitions occurring during the light period (±5.56); t *(5)* = 2.14, *p* = 0.04]”.^21^ Compared to this analysis, scores of the same data performed by SDN would have provided essentially the same results (**Figure 6 D-F**) [means (standard error (SEM): BL = 2.63 (0.24), α3KD = 3.19 (0.23); 20% change for all transitions occurring during the light period] and led to the same conclusions supported by statistical significance (one-tailed paired t-test: *t* (5) = 6.6, *p* = 0.0006).

## DISCUSSION

In this paper we describe and validate a novel transfer learning-based approach to automate and greatly accelerate the sleep-wake scoring of mouse electrophysiological data by a trained scorer. Unlike previous methods for automated scoring of rodent sleep-wake data, SDN creates a bespoke retrained model tailored to each EEG/EMG or LFP/EMG record to complete the scoring based on the end-user’s scoring style. This method is highly accurate, reliable and versatile for different experimental conditions. Accuracy is further improved by our novel REM scoring correction procedures.

Each retrained SDN model is based on GoogleNet, a deep convolution neural net (DCNN) capable of computer vision, with fine-tuning of the high-level layers to perform the classification of wake, NREM & REM, which it learns from a subset of scores from that particular file, provided by the end-user. Using transfer learning on a DCNN that was trained on images for object recognition^18^, rather than previously recorded mouse electrophysiology data means there is no bias from familiarity with older data from a specific laboratory. We believe this approach is advantageous because it adopts a scoring style from what it learns from the end-user, rather than us – the investigators who developed it. Manual sleep scoring may be regarded as the most dependable and/or the ground-truth. Thus, we argue the importance of an automated scoring method that learns to score from the end-user, rather than the investigators who developed the algorithm.

It is informative to compare our method with previously published automated sleep-wake scoring methods for rodent data. Other models work by training the model on an existing set of data and scores and using the model to score new data. Most of these are freely available. One example is “SPINDLE”^13^, a web-based service. the nature of the web-based service may arguably create difficulties depending on the institutional policies surrounding sharing of raw unpublished data. For some investigators, the ability to obtain source code and run it on a local machine, as we provide, may be an attractive alternative. Another example is “MC-SleepNet”^16^, one of its major advantages over some others like it, is the retention of time-domain information. Others extract features from Fourier transformed power spectra of epochs. This can be problematic because large amplitude, yet short, sections of an epoch may have greater representation in the power spectrum than the longer, lower amplitude, section of the epoch. This scenario poorly mimics manual scoring. SDN also maintains time-domain information by performing wavelet transforms rather than Fourier transforms. One limitation of MC-SleepNet, is that it scores 20 second epochs. In our group and others, 4 second epochs is typically used and considered necessary to capture the fast state-transitions present in mouse sleep data. “WaveSleepNet”^12^ is a Matlab based algorithm that also employs transfer learning of GoogleNet. However, unlike SDN which generates novel models tailored to individual EEG/EMG or LFP/EMG records, WaveSleepNet is a single model that users would deploy to score new data, as with the other models discussed here, it is familiar with the developers’ data sets. The available code we found for WaveSleepNet did not process the raw signals into images that WaveSleepNet would require for classifying. This may introduce challenges for investigators interested in using WaveSleepNet for their data. Additionally, we report considerably higher overall F1 scores – 0.82 by WaveSleepNet vs 0.96 by SDN. IntelliSleepScorer^15^ is another example of the more commonly found model that is trained on existing mouse EEG, expected to perform well on new data. IntelliSleepScorer provides 10 second epochs only whereas SDN does not require a predetermined epoch duration. SDN handles any epoch duration automatically, notwithstanding our post-hoc adjustments were tested in 4 second epochs only. IntelliSleepScorer also assumes signals are EEG from a frontal screw electrode and a parietal screw electrode requiring stringent channel designations. SDN does not require this. Reliability metrics of SDN were slightly higher than IntelliSleepScorer’s metrics, but considerably higher than those on data from an outside laboratory. The paper reported precision and recall separately, so we calculated F1 scores from their tables (Wang et al., 2023). For REM, IntelliSleepScorer had an F1 score of 0.72 vs 0.86 by SDN. As they report, F1 score of 0.86 is consistent with two human scorers. For NREM, IntelliSleepScorer had an F1 score of 0.89 vs 0.95 by SDN. For wakefulness, IntelliSleepScorer had an F1 score of 0.92 vs 0.97 by SDN.

One crucial consideration with most models trained with existing mouse EEG and scores is the question of whether they will perform as well as in their source publication, when presented with novel data.^17^ Both the WaveSleepNet^12^ and IntelliSleepScorer^15^ papers address this by obtaining outside data sets. In both papers, their models perform worse with outside data than with their in-house data. Presumably, this is because the model is simply more familiar with signals and scores from the same laboratory as signals and scores used to train it in the first place. Our method has the advantage that the lower layers are not trained on mouse EEG/LFP, and the higher levels are trained on signals from the individual file being scored. Thus, when shown new data, the model is no less familiar with new data than data we used in this manuscript to validate the method. We have included data generated independently by different investigators and scored by those different investigators, including data using EEG and LFP signals. Despite these differences in recording method, we found similar reliability metrics. This presents a fundamental difference with other published machine learning based sleep scoring algorithms that we have found.

Somnivore is proprietary software that reportedly works very well. It is the one exception we know of that may partially circumvent the issue of familiarity with data on which it was trained, by retraining its network on a subset of manually scored epochs from each file.^36^ The architecture of this model is not described in the publication^36^, so it is difficult to ascertain whether the model is trained on time-domain or frequency domain information, or whether the base model is originally trained on existing EEG data sets and manual scores available to those who developed it, unlike SDN which has no bias from familiarity with existing electrophysiologic data from specific laboratories. Moreover, since Somnivore is proprietary, it may not be available to many investigators who could benefit from a highly accurate sleep-wake algorithm. The reliability of SDN is in a similar range to Somnivore. However, Somnivore reports poorer accuracy at transitions. We have found our method deals with transitions well, as shown in our analysis of NREM-REM transitions. We feel this is an important attribute to conserve in sleep-wake scoring since we routinely analyze power dynamics at state transitions and bout analyses. Additionally, Somnivore reported a loss in accuracy when scoring drug-induced NREM, whereas SDN performed well when scoring drug-induced NREM. SDN users do need access to Matlab, but it is otherwise freely available. Additionally, SDN classified wakefulness and NREM slightly better than Somnivore with rodent data. Somnivore is reportedly extremely fast. We have not tested this firsthand because we have not purchased the software. SDN completed the scoring of one 24-hr file in about 10 min using the machine specifications in our methods section.

Finally, one of the major advantages of SDN is its ability to identify files that yield the lowest score-reliability. Our post-hoc modification provides this information in the absence of a set of manual scores to serve as the ground-truth, the existence of which would negate the need for an automated method. In other publications, especially for REM, we see long distribution tails into low reliability metrics. In other words, while the average reliability of a data set may be arguably acceptable, there are typically some files that yield far poorer REM reliability.^15^ Other methods do not have a means of flagging these unreliable scores. Investigators using other methods would need to accept that an unknown subset of the files in their data set were unreliable. Our method enables investigators to remove those files, or include them after scoring them manually. This would be advantageous to investigators who require the scores included in their data set to be reliable.

Who is this method for? Our aim in developing this method was to provide a tool that will take the burden of manual sleep scoring away from investigators who would otherwise prefer to score their data manually. One potential concern with using automated sleep-wake scoring methods, including machine learning based ones, is that the automated scores may poorly match the investigator’s scores. In principle, supervised machine learning based approaches should score more “like” humans because they are trained on epochs labeled by human experts. However, previously published models were trained on already existing labeled epochs scored manually by humans. It is not clear if one of these models would score similar to a new investigator using the model for the first time. Indeed, different investigators do not consistently score manually like one another.^15^ Thus, a user would need to be reasonably certain that their manual scoring style is similar to those who labeled the epochs used to train the model they are using. Our aim was to develop a method that will score “like” the investigator who is using it. Thus, our transfer learning-based approach re-trains a bespoke model for each EEG/EMG or LFP/EMG record it is given, based on a subset of epochs scored by that investigator. It is not trained on, or familiar with, epochs labeled by other individuals. Additionally, users need enough experience with manual sleep scoring that they would ensure all the states are well represented. For instance, scoring the first two-hours of the dark period would be a mistake because NREM and REM would be poorly represented. REM is of particular concern because it is underrepresented physiologically compared to wake and NREM. We manually scored a 2-hour period from ZT 2.5-4.5, a region that appeared to have REM well represented. Who is this method not for? Since our method requires a subset of manually scored epochs, it is likely not useful to anyone who is not well versed in sleep-wake scoring. An investigator who requires an algorithm to score their data for them because they do not have experience scoring manually will need to use one of the other methods, and we recommend one of those named above. Finally, we propose the secondary optional post-hoc REM correction we term ‘rule_123’ is used by investigators who have existing data sets they have scored manually to test this option first. We suggest the appropriate settings are determined by comparisons with their existing manual scores before using rule_123 in new data.

The aim of this method is to reduce the burden of manual scoring without sacrificing the scoring approach employed by an individual expert scorer. Estimates of the burden of sleep scoring from expert scores suggest a single 24-hour record requires about two-six focused labor-hours to complete. However, this cannot simply be multiplied by the number of records to score to obtain the total number of hours needed to complete a data set. Manual scoring requires concentration and leads to fatigue. Expert scorers in our labs report being able to score a single 24-hour EEG/LFP/EMG record per day, maximum. More commonly, a 24-hour record is completed gradually in separate sessions in between other work requirements. This would indicate a data set of 11 records would require, at a minimum, more than a week to complete. Manually scoring approximately 2-hours worth of data can be completed in about 10 minutes, amounting to roughly 2 hours for a batch of 11 records. 11 records then takes approximately 2 hours to complete with SDN, which takes place unsupervised by the computer, such as overnight. This indicates a burden reduction from one-two weeks of intermittent human labor to under 4 hours total time, with 80% of that time being completed without the investigator. Many pre-clinical datasets are much larger than this. For instance, consider a study with 4 groups and 10 mice per group. Using the same estimates given above, these forty 24-hr recordings would require 2 months to score manually versus ∼1 work day of investigator time, and less than one full day of total time using SDN!

This is the first method we know of to 1) create a new custom model for each file based on transfer learning of the signal of each file and scores by the end-user to adopt the scoring style of that end-user without bias of previous familiarity with signals and scores from the developers’ lab(s). 2) Flag unreliably scored files in the absence of F1 values, which would be needed for a new data set not previously scored manually. 3) validate reliable performance when scoring hypnotic-drug induced sleep compared with non-induced sleep. 4) validate reliable performance when scoring using LFP data. Given the recent boom in machine learning applications for automating sleep scoring in mouse models, the advantages and disadvantages of our method are discussed in the context of the others we know to be available.

In summary, we provide the basic and translational sleep research community with an automated sleep-wake scoring algorithm that will adopt the individual investigators manual scoring style, while reporting the files on which it could not perform well. We hope this will reduce the bottleneck in the work flow from data collection to novel findings from mouse models used to advance sleep research.

## Author contributions

FK: Conceptualization, Data curation, Formal analysis, Funding acquisition, Investigation, Methodology, Project administration, Resources, Software, Validation, Visualization, Writing – original draft, Writing – review & editing. TJS: Conceptualization Writing – original draft, Writing – review & editing. REB: Conceptualization, Funding acquisition, Project administration, Resources, Supervision, Writing – original draft, Writing – review & editing. RB: Conceptualization, Funding acquisition, Project administration, Resources, Supervision, Writing – original draft, Writing – review & editing. DSU: Conceptualization, Data curation, Formal analysis, Funding acquisition, Investigation, Methodology, Project administration, Resources, Software, Supervision, Validation, Visualization, Writing – original draft, Writing – review & editing.

---

**Figure S1.**
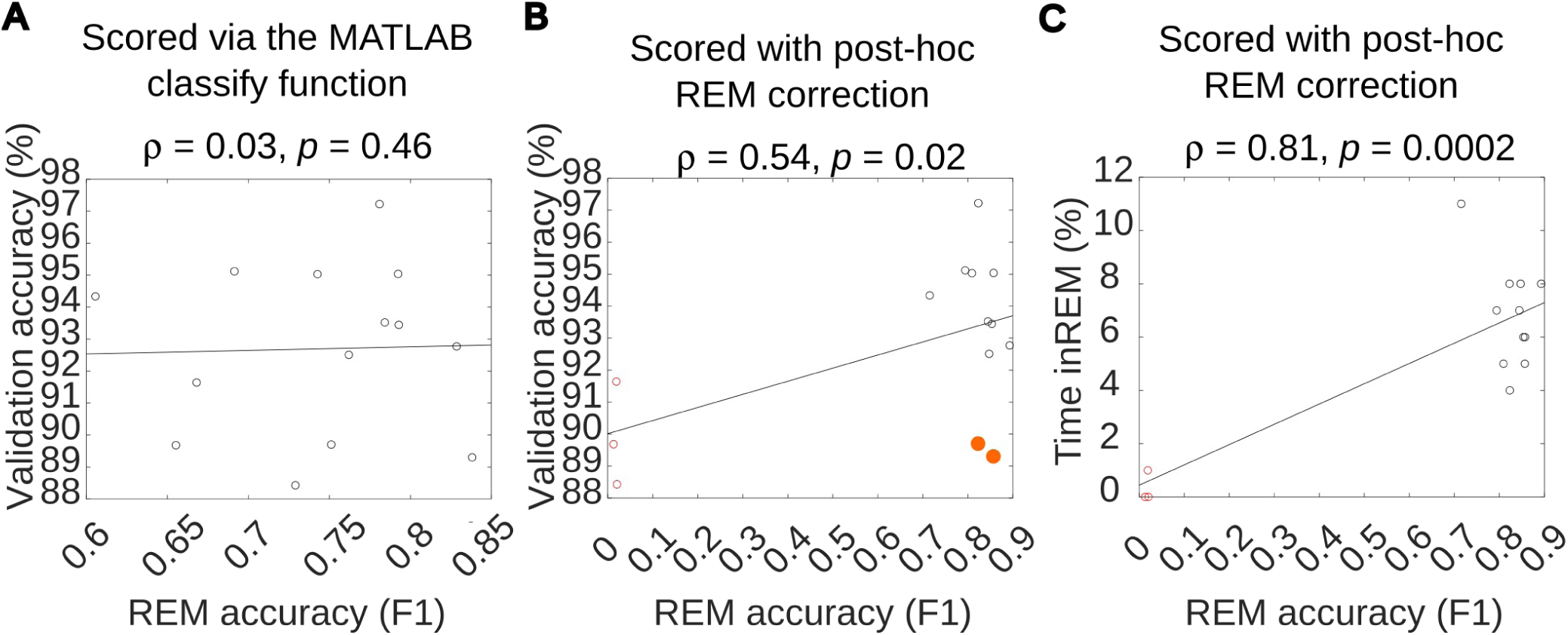
Post-hoc correction of SDN generated REM epoch scores removes unreliably scored files. **A.** Reliability of REM scores is poorly predicted by validation accuracy when REM epochs are labeled by the MATLAB classify function. **B.** Reliability of REM scores is well predicted by validation accuracy when REM epochs are corrected by our post-hoc adjustment. **C.** Post-hoc correction of REM epochs reveals files that were scored unreliably by removing REM in those files. Data were from fourteen 24-hour recordings from seven mice.

**Figure S2.**
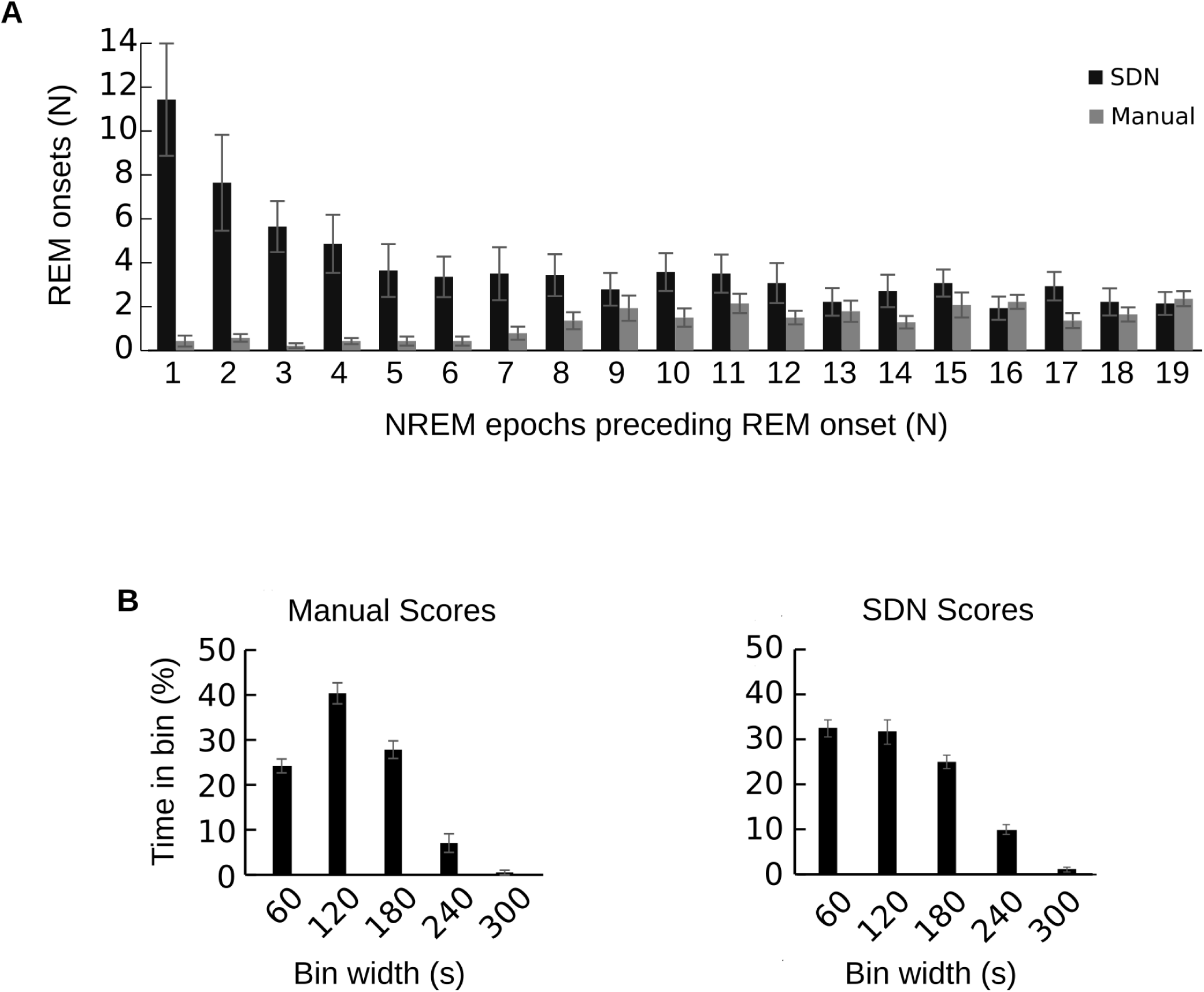
REM bouts preceded by short NREM bouts were a major source of false positive REM scores by SDN before rule_123 was deployed and poorly recreated the time-weighted bout profile for REM. **A.** The number of REM onsets preceded by different numbers of NREM epochs revealed SDN scored many REM epochs following very short NREM bouts, whereas manual scoring very rarely finds REM after short NREM bouts suggesting SDN erroneously scores early light NREM as REM. SDN identified REM onsets preceded by longer NREM bouts in a much more similar amount to manual scoring. **B.** Without rule_123, SDN overestimated REM in the shortest time-weighted bout bin at the expense of the second shortest time-weighted bout bin. Data were from eleven 24-hour recordings from six mice.

